# Mathematical model of chromosomal dynamics during DNA double strand break repair in budding yeast

**DOI:** 10.1101/2022.02.01.478611

**Authors:** Shinjiro Nakahata, Masashi Fujii, Akinori Awazu

**Affiliations:** Graduate School of Integrated Sciences for Life, Hiroshima University, Hiroshima, Japan; Research Center for the Mathematics on Chromatin Live Dynamics, Hiroshima University, Hiroshima 739-8526, Japan

## Abstract

During the repair of double-strand breaks (DSBs) in DNA, active mobilizations for conformational changes in chromosomes have been widely observed in eukaryotes, from yeast to animal and plant cells. DSB-damaged loci in the yeast genome showed increased mobility and relocation to the nuclear periphery. However, the driving forces behind DSB-induced chromatin dynamics remain unclear. In this study, mathematical models of normal and DSB-damaged yeast chromosomes were developed to simulate their structural dynamics. The effects of histone degradation in the whole nucleus and the change in the physical properties of damaged loci due to the binding of SUMOylated repair proteins were considered in the model of DSB-induced chromosomes based on recent experimental results. The simulation results reproduced DSB-induced changes to structural and dynamical features by which the combination of whole nuclear histone degradation and the rigid structure formation of repair protein accumulations on damaged loci were suggested to be primary contributors to the process by which damaged loci are relocated to the nuclear periphery.

## Introduction

Genomic DNA is frequently damaged by endogenous and exogenous factors [1]. Double-strand breaks (DSBs) are among the most serious types of DNA damage; unrepaired DSBs may lead to deletions or translocations of genome sequences that induce cancer [2-4]. Therefore, eukaryotes have evolved the ability to recognize and repair DNA damage.

Recent studies have suggested that molecular machineries and processes, such as DNA repair proteins and their SUMOylation, are widely conserved from simple unicellular yeast to higher multicellular organisms [5-12]. Additionally, during the repair process, an increase in the mobility of damaged loci and changes in chromosome conformations have also been observed as widely conserved DNA damage responses (DDRs) among eukaryotes [12-18].

Studies on DDRs in budding yeast have aided in developing a model system for eukaryote DDRs. In the yeast genome, the loci damaged by DSBs exhibited increased mobility [13-15] and relocation to the nuclear periphery [19]. Moreover, global histone degradation was observed according to DSB, which was expected to induce decomposition and contribute to changes in the mobility of damaged chromatin [20]. However, the driving forces behind the changes in chromosome conformations and the relocation of damaged loci remain unclear.

In this study, we developed models to represent the intranuclear dynamics of normal chromosomes (named normal model) and chromosomes with DSB-induced damage and histone degradation (named DSB model) in budding yeast. Simulations using these models aimed to reveal the driving forces and physical mechanisms of the DSB-induced increase in chromatin mobility and relocation of damaged loci to the nuclear periphery. Recently, several intranuclear chromosome models of normal budding yeast have been proposed [21-25]. In this study, a more simplified model of normal yeast chromosome than recently proposed ones was considered, by which the primary mechanism of chromosomal dynamics during the repair of DSBs in DNA could be revealed.

## Materials and Methods

### Coarse-graining Models of Local Structures of Normal and DSB-induced Damaged Budding Yeast Chromosomes

Coarse-grained models of normal and damaged chromosomes in the budding yeast nucleus, named the normal and DSB models, respectively, were developed as follows. The *i*-th of 16 chromosomes was described as a chain consisting of *N*_*n*_ particles with excluded volumes connected by a spring. Here, *N*_*n*_ was assumed to be proportional to the number of base pairs in the *n*-th chromosome, other than the rDNA region (Table 1). Particles in *i*-th chain were indexed by *i*= 1,2,3,…,*N*_*n*_ from the upper to lower stream of the DNA sequence of the nth chromosome. Each particle was assumed to describe a 1 kbps DNA region for both normal and DSB models and to contain five sets of nucleosomes involving 150 bps DNA with a linker involving 50 bps DNA in a normal model. A recent experimental study showed that 20 − 40% of histones degraded uniformly throughout the nucleus in response to DSB damage [20]. Therefore, in the DSB model, each particle was assumed to contain three sets of nucleosomes with a linker of 550/3 bp on average.

**Table 1.**
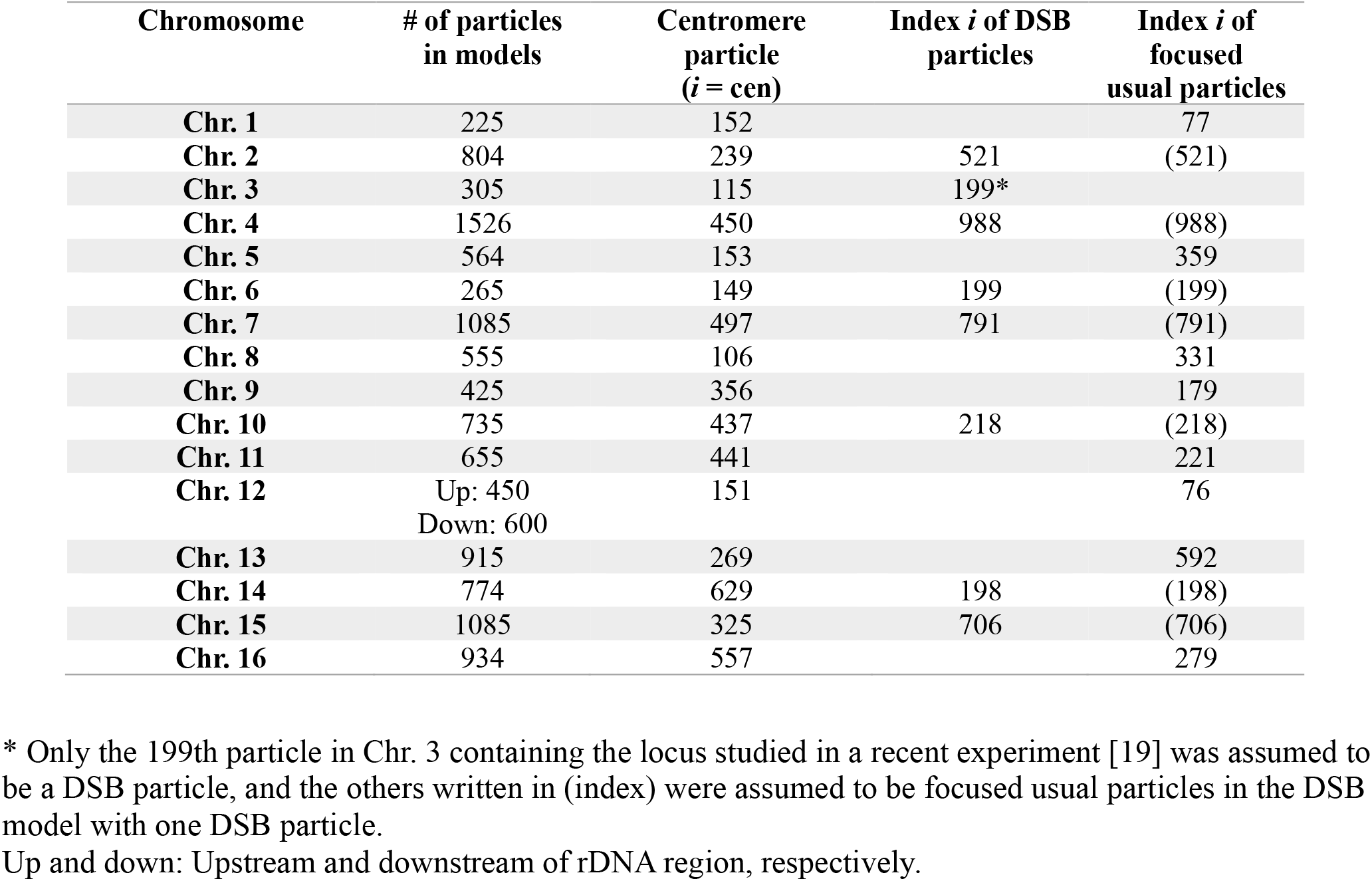
Numbers and indices of particles containing specific genome regions in each chromosome

| Chromosome | # of particles in models | Centromere particle ($i = \text{cen}$) | Index $i$ of DSB particles | Index $i$ of focused usual particles |
| --- | --- | --- | --- | --- |
| Chr. 1 | 225 | 152 |  | 77 |
| Chr. 2 | 804 | 239 | 521 | (521) |
| Chr. 3 | 305 | 115 | 199* |  |
| Chr. 4 | 1526 | 450 | 988 | (988) |
| Chr. 5 | 564 | 153 |  | 359 |
| Chr. 6 | 265 | 149 | 199 | (199) |
| Chr. 7 | 1085 | 497 | 791 | (791) |
| Chr. 8 | 555 | 106 |  | 331 |
| Chr. 9 | 425 | 356 |  | 179 |
| Chr. 10 | 735 | 437 | 218 | (218) |
| Chr. 11 | 655 | 441 |  | 221 |
| Chr. 12 | Up: 450<br>Down: 600 | 151 |  | 76 |
| Chr. 13 | 915 | 269 |  | 592 |
| Chr. 14 | 774 | 629 | 198 | (198) |
| Chr. 15 | 1085 | 325 | 706 | (706) |
| Chr. 16 | 934 | 557 |  | 279 |

\* Only the 199th particle in Chr. 3 containing the locus studied in a recent experiment [19] was assumed to be a DSB particle, and the others written in (index) were assumed to be focused usual particles in the DSB model with one DSB particle.
Up and down: Upstream and downstream of rDNA region, respectively.

The positioning and orientation of nucleosomes and linkers in chromosomes are expected to involve randomness. Thus, the radius of the *i*-th particle in the *n*-th chain, *r*_*i,n*_, was assumed to be proportional to the edge-to-edge distance of 1 kbps DNA with *m* sets of nucleosomes with linker DNA, estimated as *α*_*i,n*_ · [Linker length] · *m*^0.6^ according to the arguments of random polymers with excluded volume. In the normal model, *α*_*i,n*_ = 0.67 was assumed for all particles by which *r*_*i,n*_∼15 *nm* was obtained, which is consistent with the recently reported width scale of chromatin fibers. In the DSB model, *α*_*i,n*_ = 0.67 was also assumed, by which *r*_*i,n*_∼ 40 *nm* was obtained for each particle. Such increases in *r*_*i,n*_ of particles in the DSB model indicated that the average inter-loci distances in chromosomes with DSB-induced increased compared to those in normal chromosomes in general, which is consistent with recently reported experimental results [26].

Note that the DSB model consists of particles describing chromatin regions with and without damaged loci, named usual particles and DSB particles, respectively. The rigidity of each particle is also expected to depend strongly on the differences in chromatin states characterized by the number of nucleosomes and DNA-binding protein accumulations. The rigidity parameter of the *i*-th particle in the *n*-th chain, *q*_*i,n*_, was estimated as follows:

Based on the conventional arguments of the rubber elasticity of polymers, for all particles in the normal model and usual particles in DSB model, 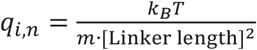was assumed if each particle describes the structure with *m* sets of a nucleosome and a linker DNA region; *q*_*i,n*_ = 9.94 × 10^−9^ *kg* · *s*^−2^ and 1.23 × 10^−9^ *kg* · *s*^−2^ were obtained for normal model and usual particle in DSB model, respectively.

Conversely, large complex accumulations of SUMOylated repair–related DNA-binding proteins, such as Mre11, Rad50, and Xrs2, are known to form on and around the damage loci [9, 12]. Therefore, for DSB particles in the DSB model, *q*_*i,n*_ was expected to be larger than that of the usual particles (Figure 1d). In the present model, *q*_*i,n*_ for each DSB particle was simply assumed to be 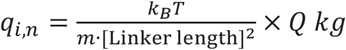 · *s*^−2^ with *Q* = 10 because there were no plausible methods for this estimation, by which *q*_*i,n*_ = 1.23 × 10^−8^ *kg* · *s*^−2^ was assumed for DSB particles. Notably, qualitatively, the same results as shown in the latter were confirmed in the case of *Q*= 3 (data not shown).

**Figure 1.**
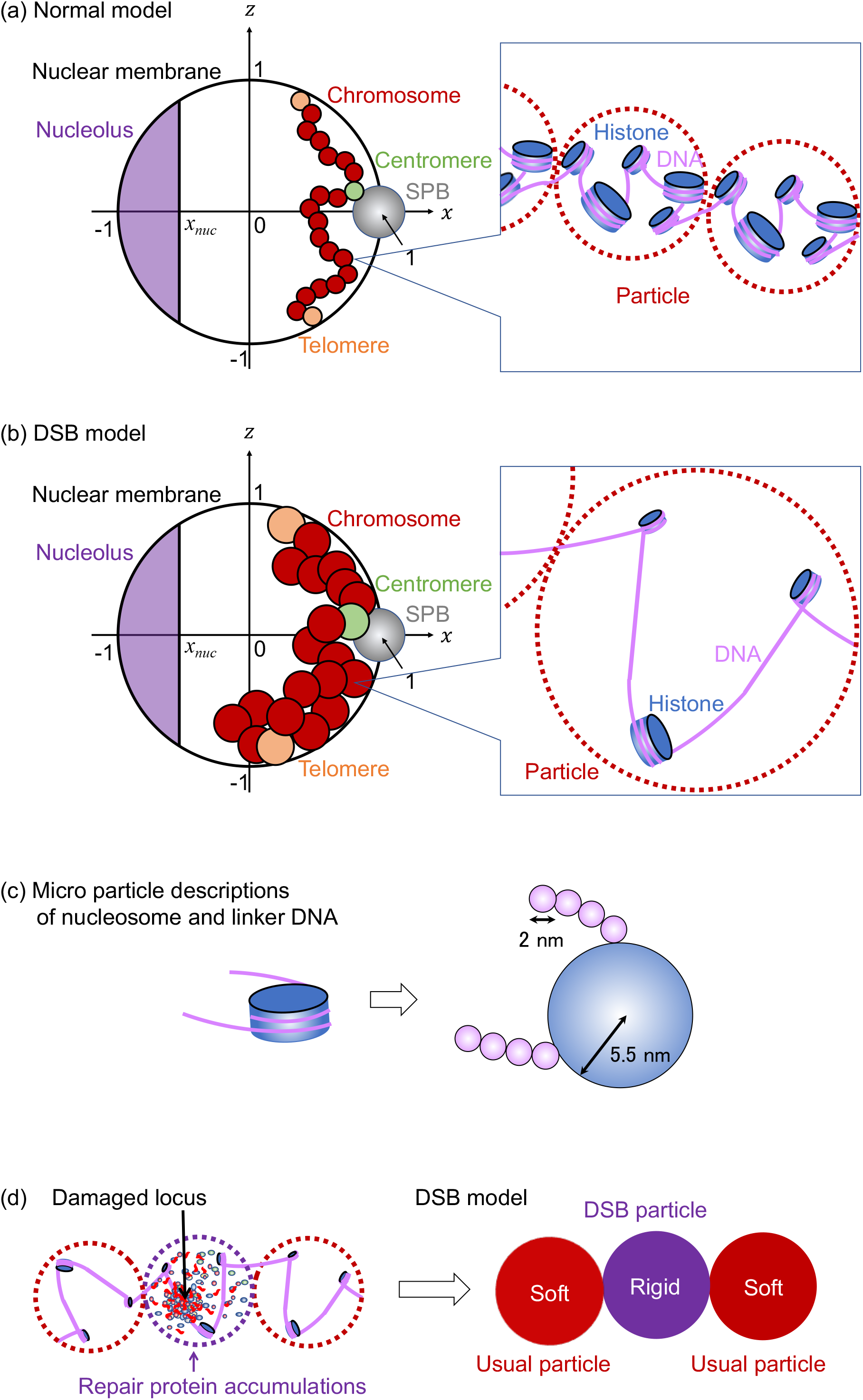
Model illustration of budding yeast chromosomes in nucleus. (a-b) Illustration of normal model (a) and double-strand break (DSB) model (b). Each particle (red) described genomic region containing 1 kbp DNA with histones. Rabble orientations were assumed where the spindle pole body (SPB) and nucleolus were positioned at the other end of the nucleus, centromere particles (green) were connected to the SPB, and telomere particles (orange) were connected to the nuclear membrane. (c) Models of nucleosome and linker DNA described by microparticles with respective radii to estimate the drag coefficient of each particle. (d) Model of DSB particles that described chromatin region containing loci damaged by DSB and large amounts of repair-related protein accumulations.

### Movement of Local Chromosome Parts

Because each part of the chromosomes was expected to exhibit 3-dimensional Brownian motion in the yeast nucleus, the position of the *i* -th particle in the *n*-th chain in the (*x, y, z*) 3-dimensional space, ***x***_*i,n*_ = (*x*_*i,n*_, *y*_*i,n*_, *z*_*i,n*_), was assumed to obey the Langevin equation, as follows:

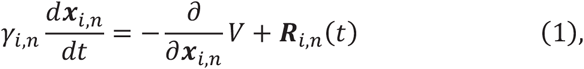

where *γ*_*i,n*_ and ***R***_*i,n*_(*t*) were coefficients of drag force and Gaussian white noise, respectively, playing the role of the random force from nucleoplasm to *i* -th particle in *n* -th chain obeying ⟨***R***_*i,n*_(*t*)⟩ = 0, and ⟨ ***R***_*i,n*_(*t*) ***R***_*i*′,*n* ′_ (*t*)⟩ = 6*γ*_*i, n*_ *k*_*B*_*Tδ*_*i i*′_ *δ*_*n n*′_*δ*(*t* − *s*) with Boltzmann constant *k*_*B*_ and temperature *T*; *k*_*B*_ *T* = 4.141947 × 10^−21^ *kg m*^2^ *s*^−2^ (*T* = 300 K). Here, *δ*_*i,j*_ indicates the Kronecker delta, and *δ*(…) indicates the Dirac delta function. V indicates that the potential of the system involves the potential of forces among the particles and the effect of boundary conditions.

The drag coefficients *γ*_*i,n*_ for all particles in the normal model and the usual particles in the DSB model were assumed to be the sum of the drag coefficients of 2 sets of a nucleosome with a linker DNA. Here, each nucleosome and each 6 bps DNA region in the linker DNA were, respectively, approximated by spheres with radii *r*_*nucleosome*_ = 5.5 *nm* and *r*_*linker*_ = 1 *nm* Hence, *γ*_*i,n*_ was assumed to be = 6*πηm*(*r*_*nucleosome*_ + *r*_*linker*_ [Number of base pairs of linker]/6) with the viscosity of the nucleoplasm *η* = 0.64 *kg m*^−1^ *s*^−1^. This estimation yielded *γ*_*i,n*_ = 8.34 × 10^−7^ kg/sec for all particles in normal model and *γ*_*i,n*_ = 1.30 × 10^−6^ kg/sec for usual particles in the DSB model.

Note that the drag coefficient for DSB particles should be assumed to be larger than that for usual particles because large amounts of repair-related proteins bind to damaged loci to form large molecular complex accumulations [9, 12]. However, its precise estimation is difficult because of the lack of experimental arguments for measuring its various physicochemical features. Hence, *γ*_*i,n*_ for DSB particles was assumed as = 2.60 × 10^−6^ kg/sec, which was simply twice the value of *γ*_*i,n*_ for usual particles in the DSB model. Notably, the results were confirmed to be quantitatively independent of *γ*_*i,n*_ for the DSB particles (data not shown).

### Interactions among Local Chromosome, Intranuclear Structure, and Nuclear Membrane Parts

In both the normal and DSB models, the nucleus was assumed to be a spherical shell with radius *R* = 1 (*μm*) containing partial regions corresponding to the nucleolus and spindle pole body (SPB). The position of the center of this container was given as (*x, y, z*) = (0, 0, 0), the region of nucleolus was assumed as *x* < *x*_*nuc*_ = −0.552, and the position of the center of SPB and its radius was given as ***x***_*SPB*_ = (*x, y, z*) = (1 (*μm*), 0, 0) and *r*_*SPB*_ = 0.1*μm*. All particles were assumed to move in a spherical shell, other than the region corresponding to the nucleolus.

The potential of system W providing the forces working on and among the particles is given as

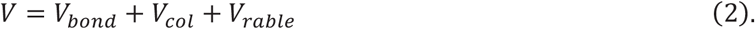

Here, *V*_*bond*_ was the interaction potential to connect each particle forming the chains corresponding to chromosomes, and *V*_*col*_ is the potential of the collisional interactions among the particles with excluded volumes, the wall of the spherical shell playing the role of the nuclear membrane, and the region corresponding to the nucleolus.

*V*_*rable*_ was the ability to form the rabble orientation of chromosomes where the centromere and telomeres of each chromosome were, respectively, associated with SPB and nuclear membrane, and a part of the 12-th chromosome, the rDNA region, was involved in the nucleolus.

The potential *V*_*bond*_ was assumed by the following:

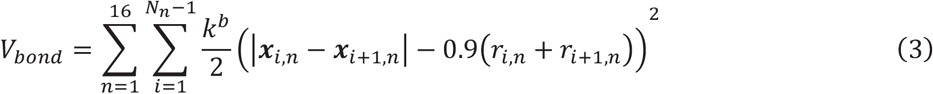

where 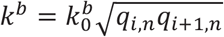is assumed to indicate the strength of the connections between neighboring particles in each chain.

The potential *V*_*rable*_ was assumed by the following:

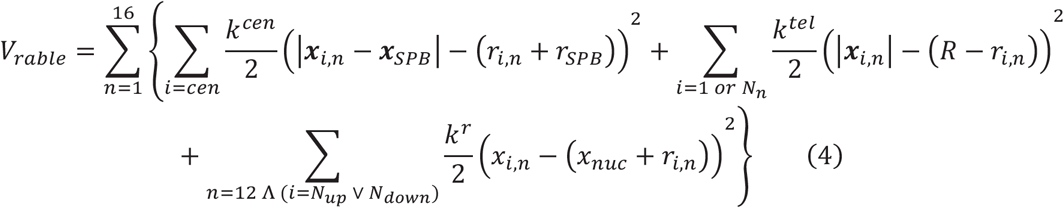

where the first and second terms indicate the potential of connective interaction between SPB and the particle describing the chromatin region containing the centromere (particle with *i* = *cen*, named centromere particle; Table 1 and Figure 1) and that between the wall corresponding to the nuclear membrane and particles describing the chromatin region containing telomeres (particles with *i* = 1 *or N*_*n*_ named telomere particles; Figure 1), respectively. The interaction strengths were assumed to be 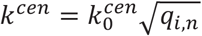and 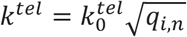were assumed. The third term indicates the potential of the connection between rDNA regions in the nucleolus and particles describing the up-and downstream neighboring regions from the rDNA region (particle with *n* = 12 Λ (*i* = *N*_*up*_ ∨ *N*_*down*_)), where 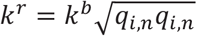was assumed.

The potential *V*_*col*_ was assumed by the following:

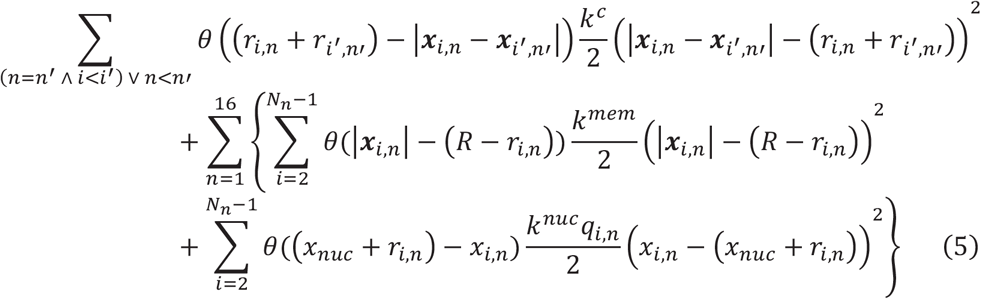

where each term indicates repulsion among particles, that between the particle and the wall corresponding to the nuclear membrane, and that between the particle and the region corresponding to the nucleolus. The interaction strengths were assumed to be 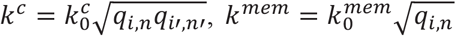, and 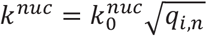 *θ* is the Heaviside step function, defined as follows:

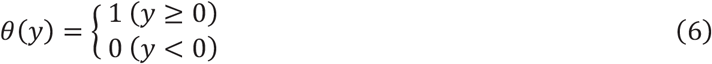

### Simulation and Statistical Analysis Methods

To simulate the model, the time integral of Langevin Eq. (1) was numerically calculated using the Euler– Maruyama method with a unit step of 0.00024 *s*. The nondimensional parameters, 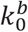and 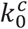, and other parameters were assumed as 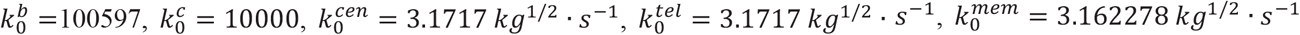, and 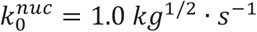in the following simulations. However, the qualitative results were confirmed to be independent of the details of these values when 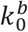and 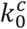were sufficiently large that the chains of particles could not pass through each other.

As the initial condition, particles were placed in the sphere along the order of the coordinates of loci inferred by 4C budding yeast experiments [27]. The mean square displacement (MSD) and radial probability density (RPD) distributions of the particles were measured using the simulation data at t= 24000− 96000 *s*, where the system appeared to relax to a steady state (Figure 2).

**Figure 2:**
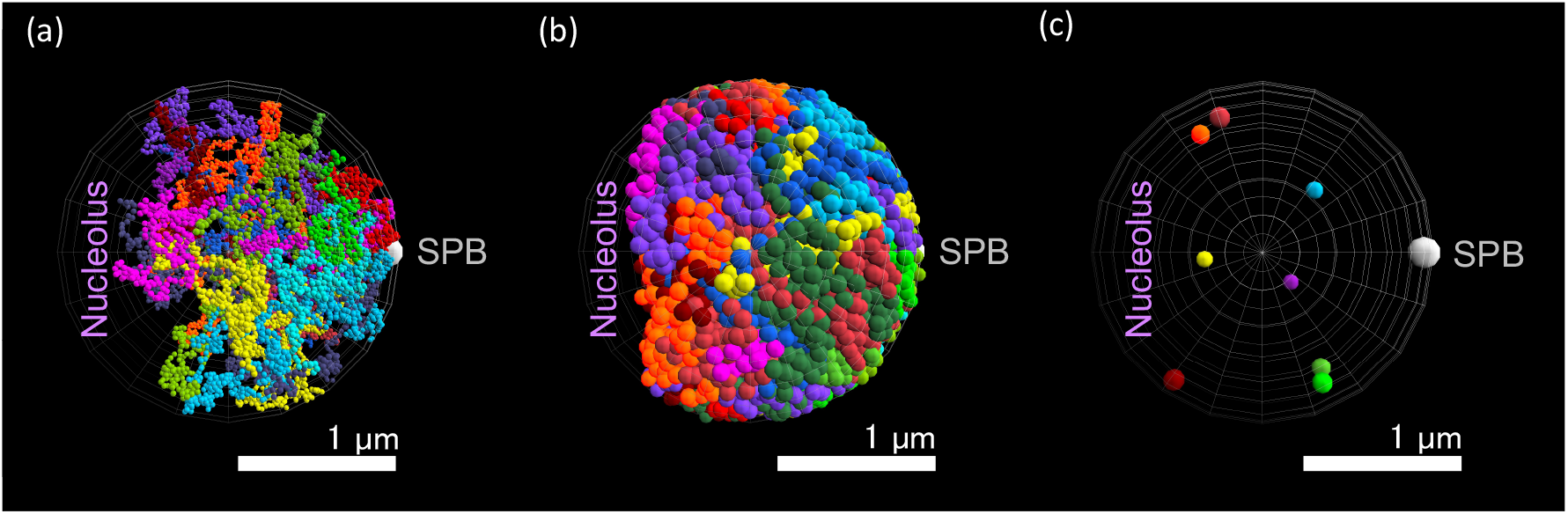
Snapshots of model simulations. (a-b) Snapshots of normal (a) and double-strand break (DSB) models; (b) simulations at 24000 *s* from initial conditions. Different chromosomes are described by chains of particles with different colors. (c) Eight DSB particles in the snapshot of simulation of the DSB model in (b).

The square displacement of the *i*-th particle was obtained by *SD*_*i*_(*τ, t*) = |***x***_*i*_(*t* + *τ*) − ***x***_*i*_(*t*)|^**2**^, MSD, and the standard deviation of SD (StdSD) of the *i*-th particle was evaluated using 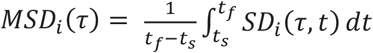 and 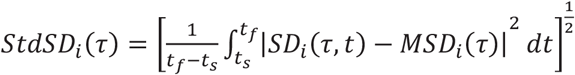with *t*_*s*_ = 24000 *sec* and *t*_*f*_ = 96000 *sec*.

The radial probability distribution of each particle is defined as the probability distribution of the distance between the centers of the particle and that of the sphere describing the nucleus. Additionally, the probability distribution of the distances between pairs of DSB particles was measured when the DSB model contained multiple DSB particles to evaluate whether the particles were scattered or clustered.

## Results

### Simulations Reproduced Increased Mobility of DSB-damaged Chromosomes

The MSDs of each particle in the normal and DSB models were measured (Figure 3). The DSB models were assumed to contain eight DSB particles (Table 1). In the DSB model, both DSB and usual particles exhibited a larger MSD than the corresponding particle in the normal model. These results were consistent with the increases in chromosome mobility reported in recent experimental studies [13-15].

**Figure 3.**
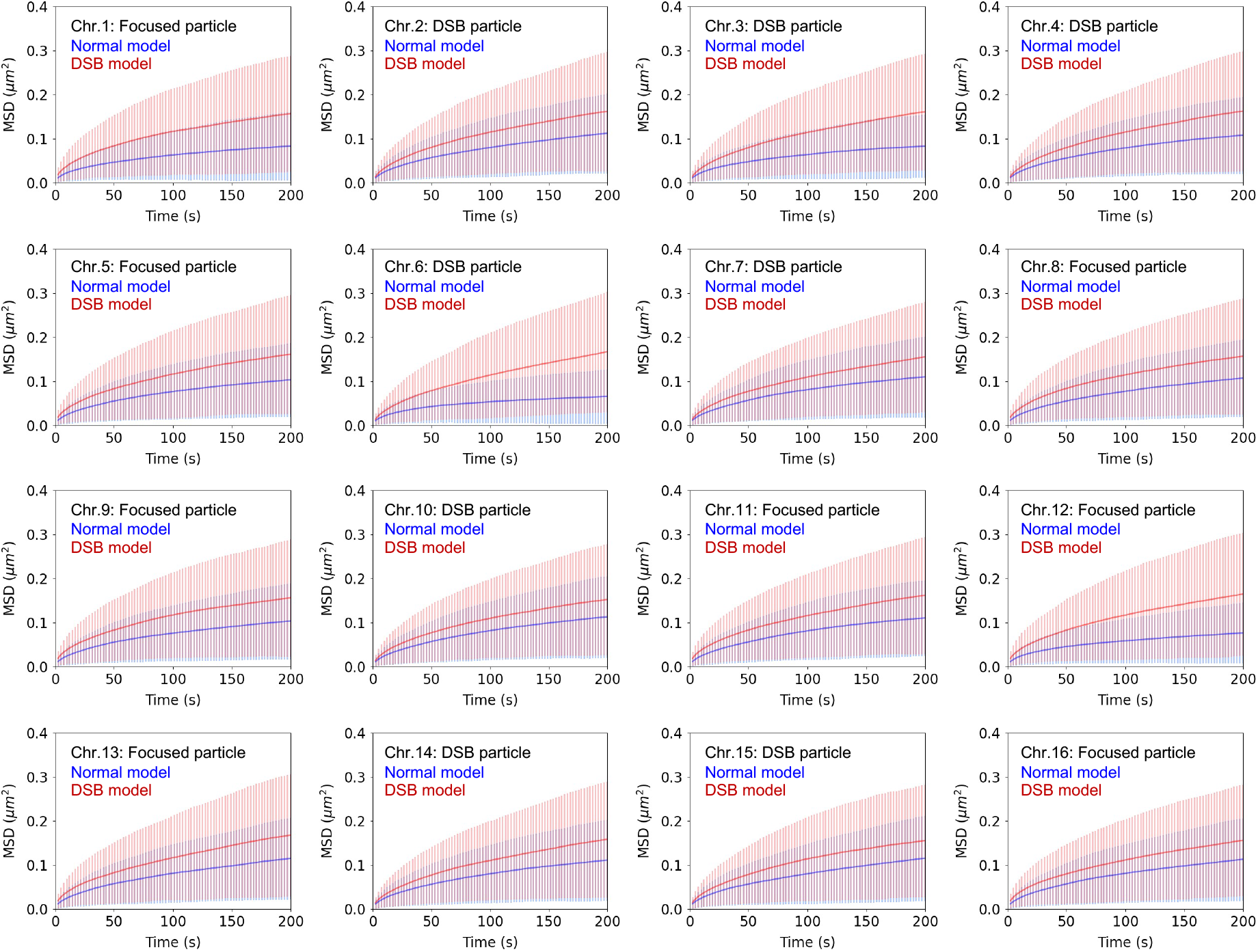
Mobility of local chromosome parts by model simulations. Mean square displacements (curves) and standard deviations of square displacements (error bars) of eight DSB particles and eight focused usual particles (Table 1) in the DSB model (red) and those of corresponding particles in the normal model (blue). Focused usual particles were selected as particles on chromosomes containing no DSB particles.

### Simulations Reproduced Relocation of Damaged Loci to the Nuclear Periphery

The RPD of the DSB particles in the DSB model and the corresponding particles in the normal model were measured (Figure 4). The DSB models were assumed to contain eight DSB particles (Table 1). The RPD of each DSB particle exhibited a steep peak near the spherical wall, playing the role of a nuclear membrane, whereas the RPD of the corresponding particle in the normal model exhibited low values. It was noted that in the DSB model, the RPD of the usual particles also exhibits large values near the spherical wall compared to those of the corresponding particles in the normal model (Figure 4). However, the RPD values of the DSB particles near the spherical wall were much larger than those of typical particles.

**Figure 4.**
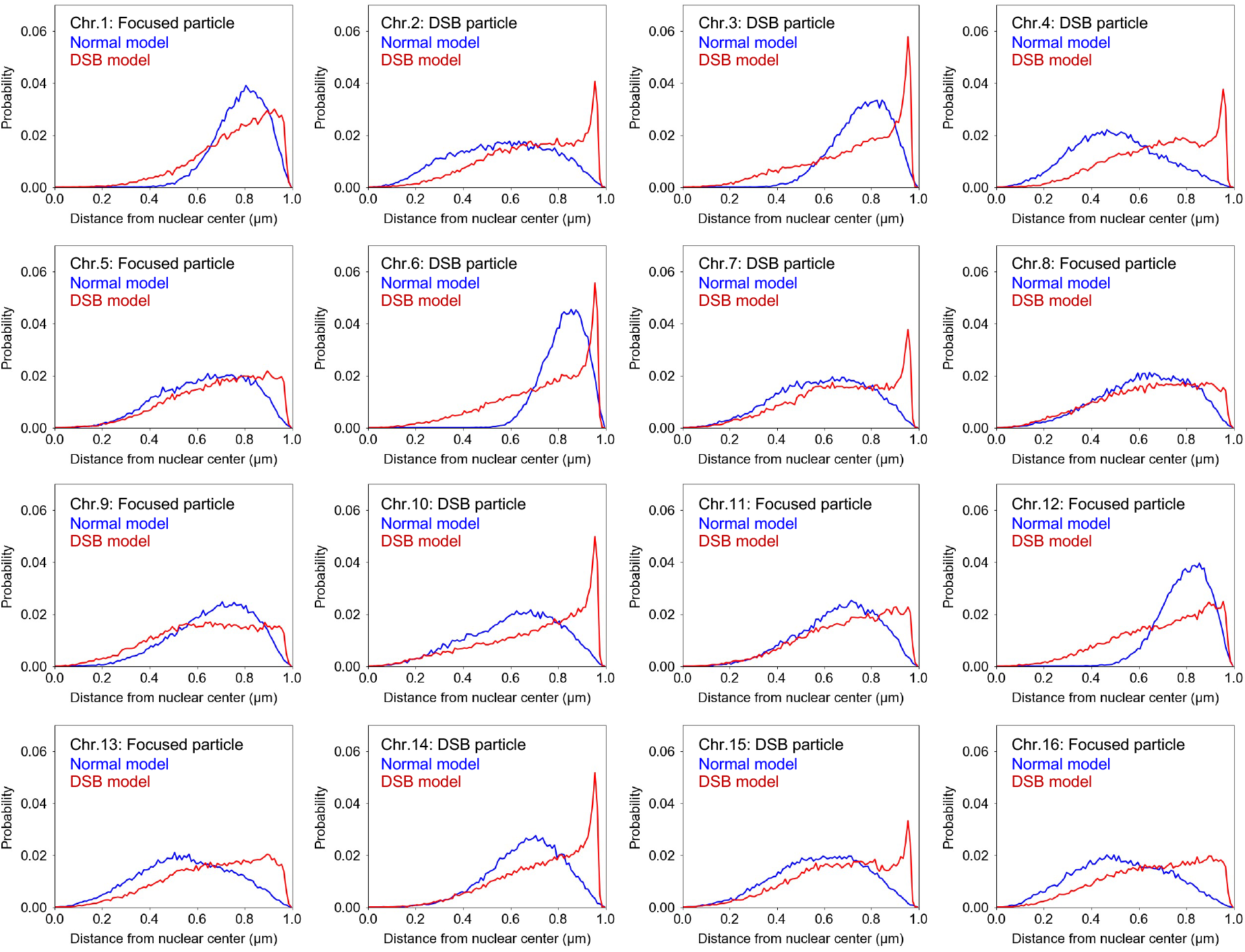
Radial probability density (RPD) of local chromosome parts determined by model simulations. RPD of eight DSB particles and eight focused usual particles (Table 1) in the DSB model (red) and that of corresponding particles in the normal model (blue). Focused usual particles were selected as particles on chromosomes containing no DSB particles.

It was noted that the same behaviors of both DSB and usual particles were obtained in both DSB models with one and eight DSB particles (Supplementary Figure S1). This indicated that the relocations of the DSB particles occur not collectively but independently.

## Discussion

The DSB model exhibited a higher mobility for each particle and specific localization of DSB particles in the vicinity of the spherical wall than the normal model. These results indicated that the present model reproduced the experimentally-observed DSB-induced phenomena in budding yeast nuclei, where the mobility of chromosomes in the whole nucleus was increased, and damaged loci were relocalized near the nuclear membrane [13-15, 19].

The length of each linker was elongated on average by the whole chromosome uniform histone degradation induced by DSBs. Following recent arguments in polymer physics, such linker elongations were considered to enlarge the volume of the effective region occupied by chromatin fibers and weaken the excluded volume effects among the effective regions. In the present study, these two effects were observed, affected by the difference in radius and rigidity of particles describing the chromatin region containing 1 kbp DNA between normal and DSB models; each particle in the DSB model was assumed to be larger and softer than that of the normal model. The present study suggests that these effects play key roles in the occurrence of DSB response behaviors in the yeast chromosomes.

The centromere of each chromosome is connected to the SPB in the yeast nucleus, which restricts the motion of the intranuclear chromosomes. However, DSB-induced expansion and softening of the effective chromatin regions weakened this restriction and extended the intranuclear mobile space of these regions. These facts could explain the increase in the mobility of chromatin regions in the whole nucleus, which is consistent with the recently proposed explanation based on observations of chromatin elongation by DSBs in the yeast nucleus [26]. It should also be noted that the simulation of the following variant of the DSB model, which consists of only usual particles with larger radii than the particles in the normal model, also shows an increase in particle mobility (Supplementary Figure S2). This suggests that the expansion of chromatin region by histone degradation is a primary contributor to the increase in chromatin mobility, which is consistent with recently reported experimental results [20].

The expansion of effective chromatin regions increases the volume fraction of chromosomes in the nucleus. Additionally, elements with different physical characteristics such as volume, shape, and rigidity tend to segregate with each other in space under crowded situations by various elements, which may be explained by the entropy effects and a similar mechanism to the depletion force [28-35] (note that the excluded volumes of rigid particles were effectively larger than those of soft particles even if the volumes of particles were uniform). Here, the chromatin regions containing DSB-induced damaged loci were expected to be more rigid than other regions due to the accumulation of DNA-binding repair–related proteins; these regions were implemented by rigid particles in the present model. These facts suggest that the chromatin region with damaged loci tends to be driven away to the nuclear periphery from the inner space of the nucleus, where non-damaged chromatin regions are crowded. In other words, the entropic effect induced by the increase in the volume fraction of effective chromosome regions in the nucleus and subsequent rigidization of the chromatin region around damaged loci was likely the driving force inducing the relocation of damaged loci to the nuclear periphery.

It should be noted that the simulation of the following variant of the DSB model, where DSB particles were assumed to be rigid but the radii of all particles remained the same as the normal model, did not exhibit any clear localization of DSB particles around the spherical wall (Supplementary Figure S3). This observation supports the importance of chromatin region expansion by histone degradation for inducing relocation of damaged loci to the nuclear periphery.

## Conclusion

Mathematical models of normal and DSB-damaged yeast chromosomes were developed and simulated. Various aspects of the mechanisms of DSB-induced chromatin dynamics in the yeast genome such as the increase in mobility of whole chromosomes and relocation of DSB-damaged loci to the nuclear periphery were revealed.

The mobility of each chromatin region in the presented simulations exhibited a much larger standard deviation than that observed in the experiments [13-15]. This is because some effects other than histone degradation might also contribute to DSB-induced increases in whole chromosome mobility, which should be discovered to prove the model in the future. The assumptions for modeling chromatin regions with damaged loci should be validated and improved from more detailed biochemical studies with experiments and analysis in the future. The clarification of the more detailed mechanism of histone degradation will also be an important future issue to describe the entire process of the DSB damage response of yeast chromosomes. Additionally, based on the progress of studies on such future issues, a model that can reproduce the damage response phenomena quantitatively will be constructed in the future.

## Author Contributions

S.N. and A.A. conceived and designed the study; S.N. and A.A. developed the mathematical model and conducted the simulations; S.N., M.F., and A.A. analyzed the data; S.N, M.F., and A.A. wrote the manuscript.

## Acknowledgements

We thank C. Horigome for providing the useful information. Computations were partially performed on the NIG supercomputer at ROIS National Institute of Genetics.

---

**Supplementary Figure S1.**
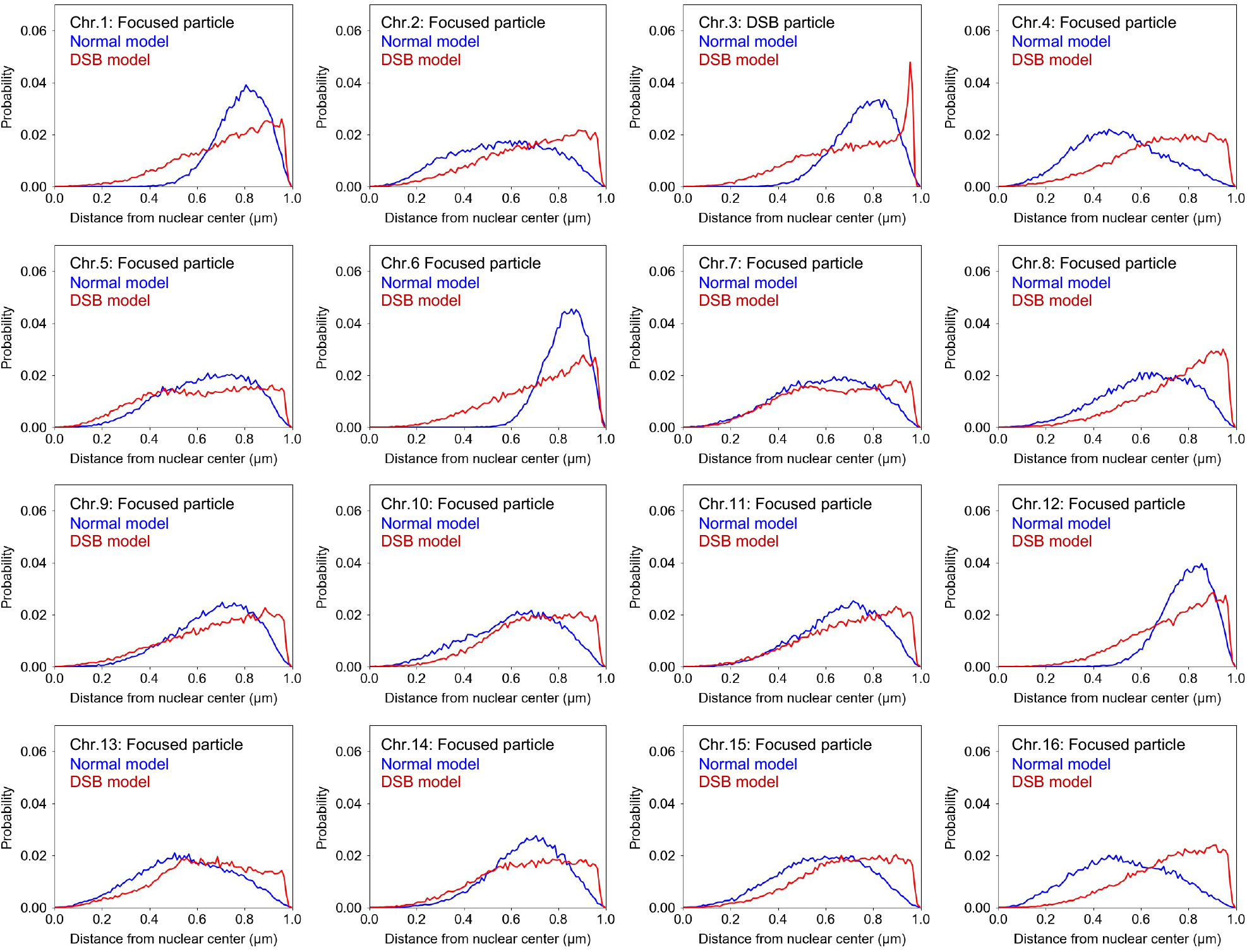
Radial probability density (RPD) of local chromosome parts by double-strand break (DSB) model simulation containing one DSB particle. RPD of one DSB particle and 15 focused usual particles (Table 1) in the DSB model with one DSB particle (red) and those of corresponding particles in normal model (blue). (b) RPD of DSB particles in the DSB model with one DSB particle (red) and that of corresponding particles in the normal model (blue).

**Supplementary Figure S2.**
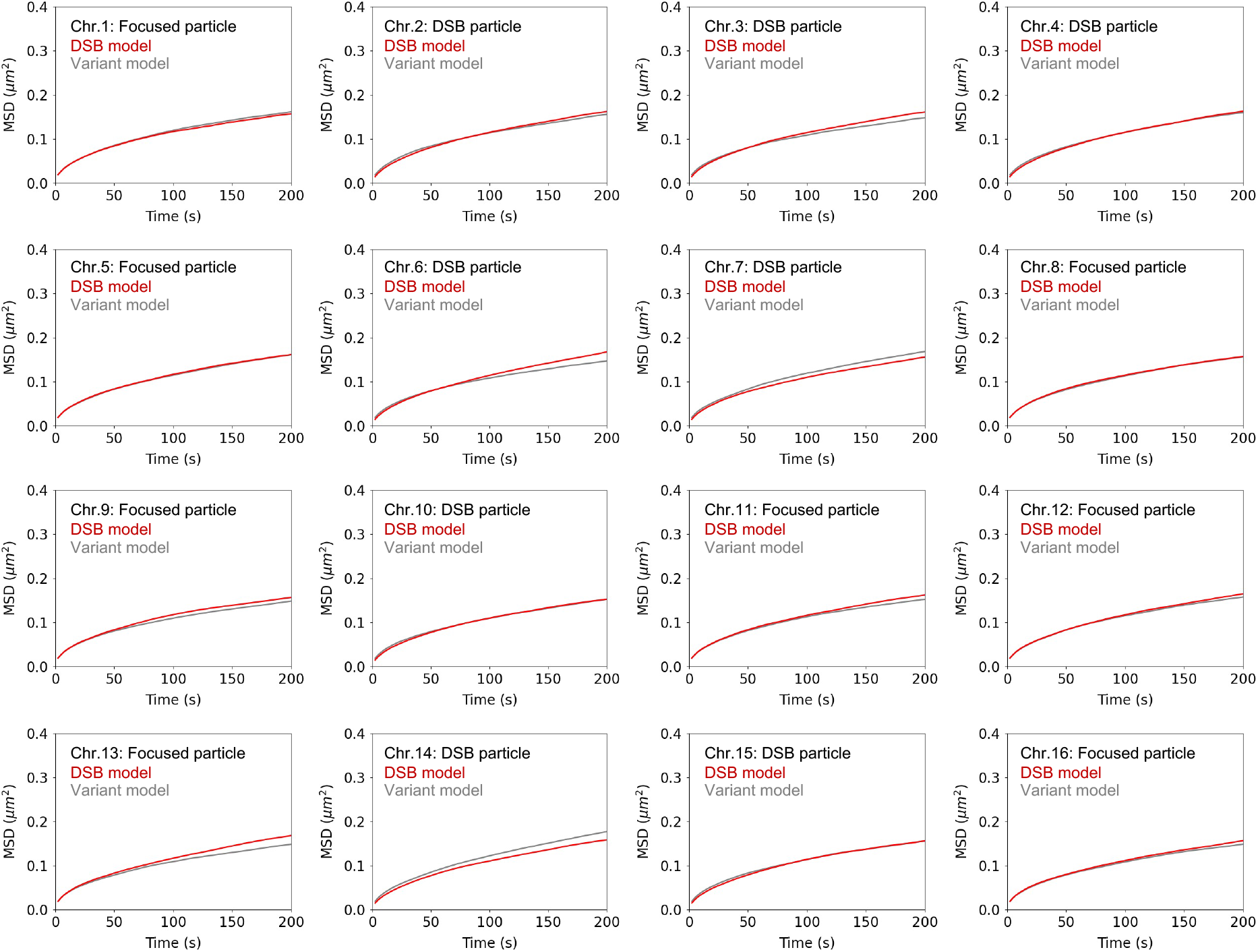
Mobilities of local chromosome parts by simulations of a DSB model variant containing no DSB particles. Mean square displacements of particles in a variant of the DSB model including only usual particles with larger radii than the particles in the normal model. Results are shown for DSB particles and focused usual particles in the DSB model (red) and the corresponding particles in the variant model (gray) (Table 1).

**Supplementary Figure S3.**
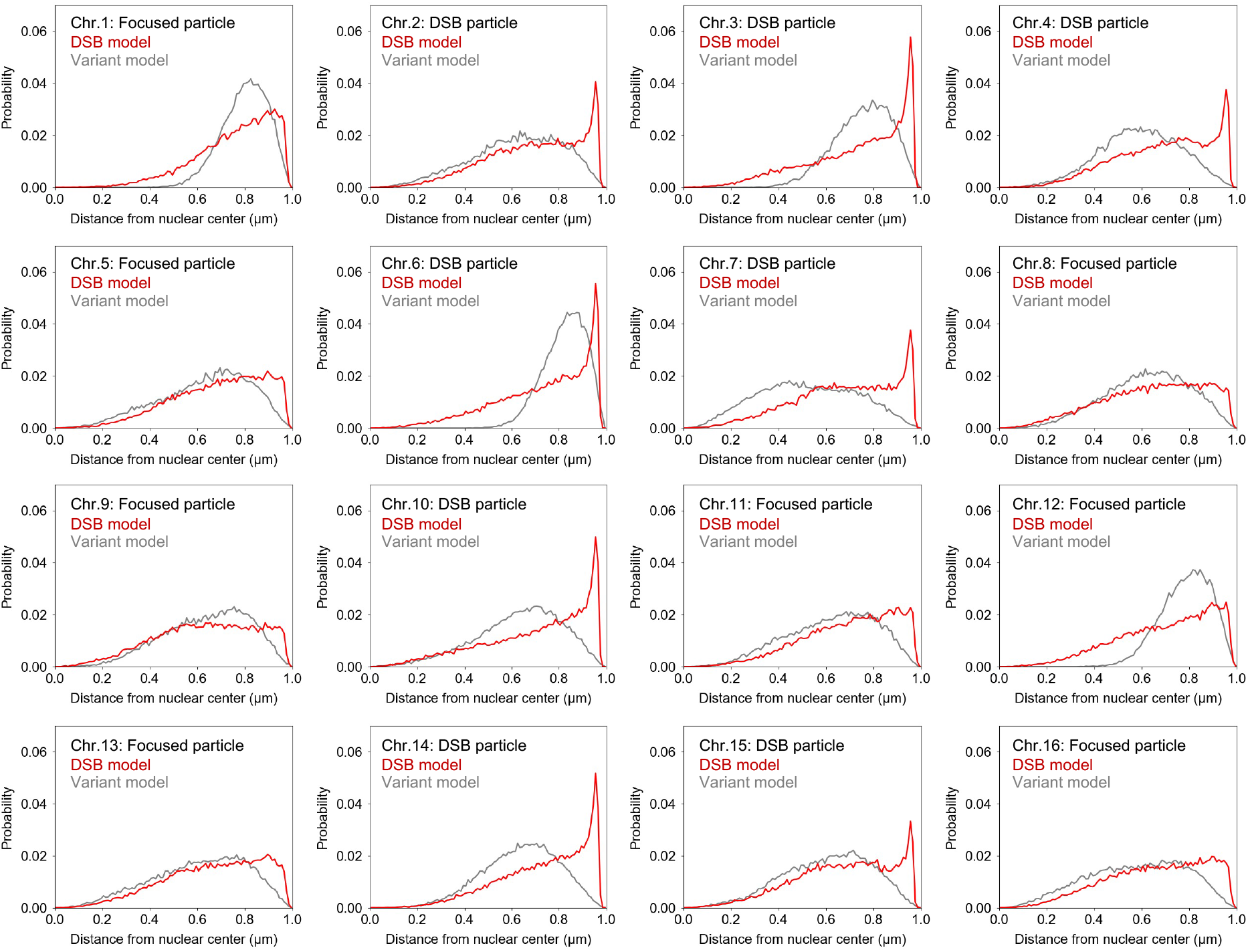
RPD of local chromosome parts by simulations of a DSB model variant in which histone degradation effects were neglected. RPD of particles in another DSB model variant in which the radii of all particles were the same as those in the normal model, but the corresponding particles to DSB particles in DSB model were assumed to be more rigid than other particles. Results were shown for DSB particles and focused usual particles in the DSB model (red) and the corresponding particles to them in the variant model (gray) (Table 1).

